# Cutting-edge microfluidic technology allows detection and sorting of single fluorescent extracellular vesicles

**DOI:** 10.64898/2026.10.06.756207

**Authors:** Mickaël Couty, Kimhuong Say, Clément Roux, Evan Lamour, Liam Barry-Carroll, Jean-Christophe Delpech, Philippe Graindorge, Beatrice Clemenceau, Sophie Bourzeix, Mathias Girault, Guillaume van Niel

## Abstract

Extracellular vesicles (EV) are cell-secreted, nanosized biological particles that are involved in most physiological and pathophysiological processes. Their composition and properties reflect the pathophysiological state of the producing cell, tissue, and organ. Released into biological fluids, they may be used as a potential source of non-invasive biomarkers for diagnosis, prognosis or therapeutic follow-up in several diseases such as cancer. However, despite the development of single-EV analysis technologies, the sorting of EV subpopulations for downstream analysis remains a major challenge. Here, we describe a novel instrument that combines a highly sensitive optical detection system with an advanced lab-on-a-chip device. Using engineered fluorescent small extracellular vesicles (sEV), we demonstrate that the Hekat NanoSorter enables sensitive, scalable, and reproducible detection and sorting of individual sEVs for downstream analysis by cryo-EM. Furthermore, detailed characterization of the sorted EV populations, together with the detection of sEVs labeled with an in vitro pathophysiological marker, demonstrates the ability of the NanoSorter to monitor and enrich clinically relevant EV-associated biomarkers. These findings highlight the clinical potential of extracellular vesicle sorting for their use as biomarkers for therapeutic monitoring and treatment outcome assessment.

## Introduction

Extracellular vesicles (EV) have been identified in nearly all organisms and cell types studied to date^1^ and have become a major field of interest across a wide range of research areas. They are now recognized as key players in cell communication and homeostasis^2–4^ in various pathophysiological processes, including cancer^5^ and are harnessed as biomarkers and drug nano-vehicles in a growing number of clinical trials^6^.

Their fundamental roles and their potential clinical applications have driven scientists to develop new technologies to ease and improve their characterization^7^. However, the study of EV presents significant biological and technical challenges. These include the overlap in their size (ranging from 30 nm to several micrometers), their density, and the heterogeneity among EV subsets, which arise from different cellular origins^8^. Such complexity often forces researchers to purify and analyze EV in bulk using isolation protocols, thereby losing critical information about the origin and individual properties of the vesicles^9,10^.

To address these challenges, new technologies, known as single vesicle technologies, have emerged. These enable refined isolation processes, advanced imaging, and more sensitive characterization, potentially revealing insights into the origin, composition, tropism, and roles of various EV subsets in physiological processes and diseases^11,12^. Despite significant progress that has improved the quality and reproducibility of EV-related studies^13^, several milestones remain unmet. In particular, the ability to efficiently and reproducibly detect and sort specific EV sub-populations from bulk samples for downstream analysis is still unresolved.

The use of flow cytometry may provide valuable answers, but it is primarily adapted for detecting and sorting cells, which are 10 to 100 times larger than EVs. Therefore, the volume of the droplets generated by the nozzles, is optimized for the sorting of cells. When applied to EVs, they limit the physical sorting and recovery of EVs and result in highly diluted samples that are unsuitable for post-isolation analysis^14^. A second limitation, that is also dependent on the droplet volume used to analyze, is the detection limit : larger droplets requires larger laser spot sizes, which causes greater photon loss, and reduced sensitivity^15^. This size discrepancy pushes cytometers to their detection limits, compelling researchers to implement stringent controls and use antibody panels. These methods are also limited by steric effects due to the small size of EV and the presence of a protein corona^16^. Nevertheless, new nano-cytometers are improving detection limits, allowing researchers to sort EV, albeit currently only through intensity gating for characterization.

Regarding droplet sorting, electric fields are commonly used to reroute droplets containing single cells from their original trajectory. However, depending on the method, the electrical impulses used in cell sorting to charge or reroute droplets can induce stress and compromise the integrity of the plasma membrane. While cells may recover from such stress, how this process affects the structural and functional properties of EV membranes post-sorting is still understudied^17,18^. Furthermore, standard flow cytometers are designed to process cells, whose size aligns with the electrostatic principles required to apply an electric field^19^. For nano-sized objects like EV, their small size introduces additional variables, such as buffer viscosity and their much lower mass, necessitating alternative approaches for effective sorting¹⁹.

To date, no instrument enables the efficient, specific, and reproducible sorting of nano-sized biological objects for the analysis of EV subpopulations post-sorting. With the ultimate goal of optimizing the therapeutic potential of extracellular vesicles, we are evaluating here a new instrument developed by Hekat Fluidics. This system combines an advanced optical detection module with a microfluidic lab-on-a-chip, capable of detecting fluorescent particles as small as 30 nm in diameter. Thanks to an ultra-thin nozzle, it generates femtoliter-scale droplets, each containing a single particle. This technology allows for the detection and sorting of small EV (sEV) with high concentration, enabling the recovery of specific, intact EV subpopulations based on fluorescence. The sorted sEV may then be used for downstream characterization, omics analysis, and functional studies.

## Results

### Engineering of CD63-fluorescently tagged sEV in a reproducible manner for NanoSorter assessment

There are already several calibration standards based on fluorescent silica or latex beads which match the size of sEV^20^. However, despite their usefulness, they do not always match the size, fluorescence and heterogeneity of biological samples which leads to a potentially high variability when they are used as standards for the analysis of biological samples. To assess Hekat sorting technology with biological samples, the first step was to generate fluorescently tagged sEV in a reproducible manner, allowing us to work with standardized biological material. To do this, we harnessed the widely used tetraspanin CD63 as an EV reporter. We first transiently overexpressed a construct of CD63 fused to mNeonGreen or mRFP fluorescent proteins at its N-terminus in HEK cells. In parallel, we pursued a long-term approach by generating stable HeLa cell lines expressing mRFP-CD63 or mNeonGreen-CD63 in a CD63 knockout (KO) background^21^.

Initial observations by confocal imaging confirmed the expression of the fluorescent reporter and its co-localization with endogenous CD63 in intracellular compartments (Supplementary Figure 1a). In stably expressing HeLa cell lines, we confirmed the expression and expected localization in endolysosomal compartments by observing its co-localization with the lysosomal marker Lamp-1 in EV-producing cells (Figure 1a).

**Figure 1.**
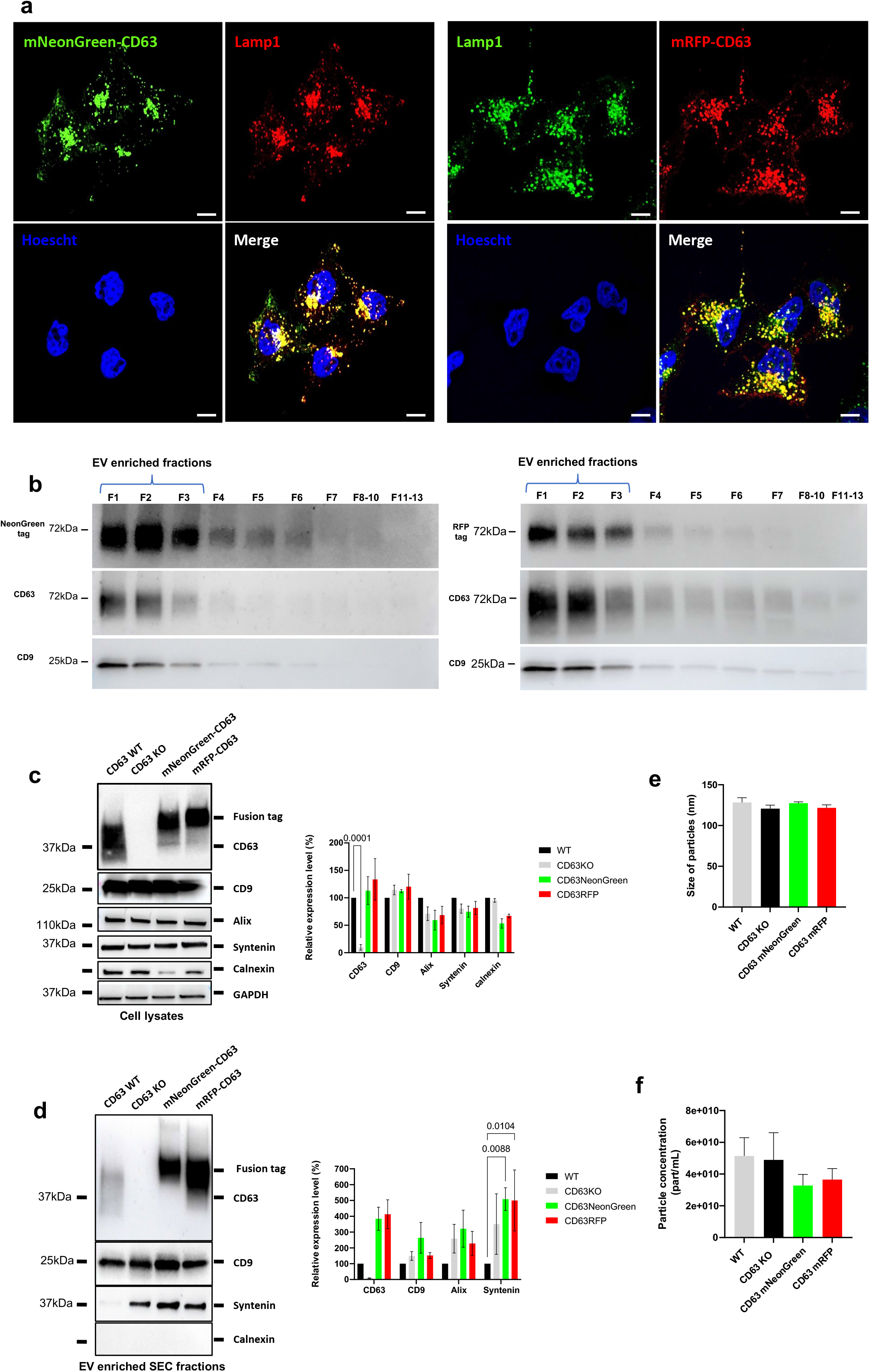
a. Subcellular localization of fluorescent reporter in HeLa mNeonGreen-CD63 and HeLa mRFP-CD63 stable cell lines. Lysosomes were labelled for Lamp1 and nucleus with Hoescht. Endogenous fluorescence for both fusion proteins was observed. Scale bar 10um. 3 independent experiments. **b.** Western blot of size exclusion chromatography (SEC) fractions from HeLa mNeonGreen-CD63 and HeLa mRFP-CD63. Fractions 8 to 10 and 11 to 13 were pooled before loading into SDS-PAGE gel wells. Samples were immunoblotted for their respective fluorescent tag along with CD63 and CD9. 3 independent experiments. **c.** Western blot of HeLa mNeonGreen-CD63, mRFP-CD63, HeLa WT and HeLa CD63 KO. Cell lysates were immunoblotted for fluorescent tags and EV markers. 3 independent experiments. Quantification shows the relative expression of each protein out of the control condition (i.e HeLa WT). 2 way Anova test. **d.** Western blot of EV enriched SEC fractions pooled from HeLa mNeonGreen-CD63, HeLa mRFP-CD63, HeLa WT and HeLa CD63 KO cells. Samples were immunoblotted for their respective fluorescent tag along with CD63 and CD9. 3 independent experiments. Quantification shows the relative expression of each protein out of the control condition (i.e HeLa WT). 2-way Anova test.; **e.** Size of HeLa mNeonGreen-CD63, HeLa mRFP-CD63 and HeLa WT derived sEV isolated by SEC. Size measured by NTA with NanoSight S300. 3 independent experiments. HeLa WT n=7, HeLa CD63 KO n=5, HeLa mNeonGreen-CD63 n= 6, HeLa mRFP-CD63 n=6; Ordinary one-way Anova test. **f.** Concentration of HeLa mNeonGreen-CD63, HeLa mRFP-CD63 and HeLa WT derived sEV isolated by SEC. Size measured by NTA with NanoSight S300. 3 independent experiments. HeLa WT n=7, HeLa CD63 KO n=5, HeLa mNeonGreen-CD63 n= 6, HeLa mRFP-CD63 n=6; Ordinary one-way Anova test.

Alongside imaging, we performed size exclusion chromatography (SEC) to isolate EV from conditioned culture media (CCM). Western blot analysis of the collected SEC fractions confirmed the expression of both mNeonGreen and mRFP fluorescent proteins in the EV fractions at a molecular weight (∼72 kDa), corresponding to their association with CD63. While we observed some cleaved forms of mRFP-CD63 (25 kDa) associated with EV from HEK cells transiently expressing CD63 constructs (Supplementary Figure 1b), we did not detect any cleavage of either fluorescent protein from CD63 in the HeLa stable cell lines (Figure 1b).

Following EV cargo characterization, we measured the size and concentration of particles isolated from CCM. Initial measurements by dynamic light scattering (DLS) or nano-flow cytometry (nFCM) revealed two different profiles for mNeonGreen-CD63 and mRFP-CD63 EV derived from transiently transfected HEK cells. We observed a larger particle diameter by DLS compared to nFCM, which correlates with observations by transmission electron microscopy. We did not observe a difference in particle concentration between the two instruments, likely due to the yield variability between EV batches (Supplementary Figure 1c–f)^22^.

As we aimed to generate reproducible EV batches to assess the NanoSorter technology, and transient expression introduced variability in EV concentration compared to HeLa stable cell lines, we used HeLa mNeonGreen-CD63 and HeLa mRFP-CD63 to produce standardized EV batches for the rest of the study.

We expanded the characterization panel of HeLa cells and their derived EV by pooling enriched EV fractions (F1-F3) and analyzing them by Western blot to detect any changes in EV-related cargo. In whole-cell lysates, apart from a molecular weight shift for CD63 due to the insertion of mNeonGreen or mRFP compared to HeLa CD63 wild-type (WT) and its depletion in CD63 KO cells—we did not detect any changes in other potential EV cargo proteins (Figure 1c). Interestingly, in EV samples, we noted a significant increase of syntenin in EV from cells overexpressing tagged-CD63 (Figure 1d). Such an increase had been previously reported in CD63 KO cells but without statistical significance^21^ and may be linked to the reported interaction of CD63 with syntenin for its integration into EV^23^. Finally, we did not observe significant changes in the size or concentration of these EV batches compared to HeLa WT controls by nanoparticle tracking analysis (NTA) confirming previous reports (Figure 1e–f). Altogether, these data led us to use HeLa mNeonGreen-CD63 and HeLa mRFP-CD63 to produce standardized EV batches to assess the NanoSorter technology.

### Microfluidic system allows detection and sorting of single fluorescent particles

Whether through light scattering or fluorescence emission, the detection of nano-sized particles is a common approach to characterize extracellular vesicle (EV) samples. Based on light scattering, Nanoparticle Tracking Analysis (NTA) is widely used by researchers. This technique relies on tracking the Brownian motion of particles to estimate their size and extrapolate their concentration in a given sample^24^. On the other hand, fluorescence-based detection methods, such as flow cytometry or nano-flow cytometry (nFCM), enable the use of specific fluorescent dyes, allowing for the targeting of particular features or markers that size alone cannot address. Combining light scattering with fluorescence detection in these methods can provide estimates of particle size and, to some extent, concentration.

To assess our standardized EV production and evaluate the detection capabilities of the NanoSorter, we established a standardized isolation and storage protocol to produce, aliquot and store multiple EV batches under identical conditions for further measurement across three different devices. First, using NTA and nFCM, we measured the mean size of our EV samples based on light scattering. NTA revealed a mean size of approximately 120 nm (±9 nm) for each EV sample, while nFCM estimated a mean size of approximately 75 nm (±4 nm) (Supplementary Figure 2a and b) confirming our previous observations (Supplementary Figure 1e and h). For both devices, no differences were observed between HeLa tagged-CD63 EV and their controls (HeLa WT and CD63 KO). Regarding particle concentration, we did not detect significant differences by NTA (1.8 × 10¹⁰ ± 0.4 × 10^10^ particles/mL) or by nFCM (1.7 × 10⁹ ± 0.6 × 10^9^ particles/mL) among the four producing cell lines, confirming the reproducibility of our EV production (Figure 2a and b). These observations, which highlight discrepancies in particle size estimates (Supplementary Figure 2c), have been previously described in the literature^22^ and emphasize the importance of defining reproducible methods for EV isolation and characterization as well as common calibration method for different instruments that use different technologies..

**Figure 2.**
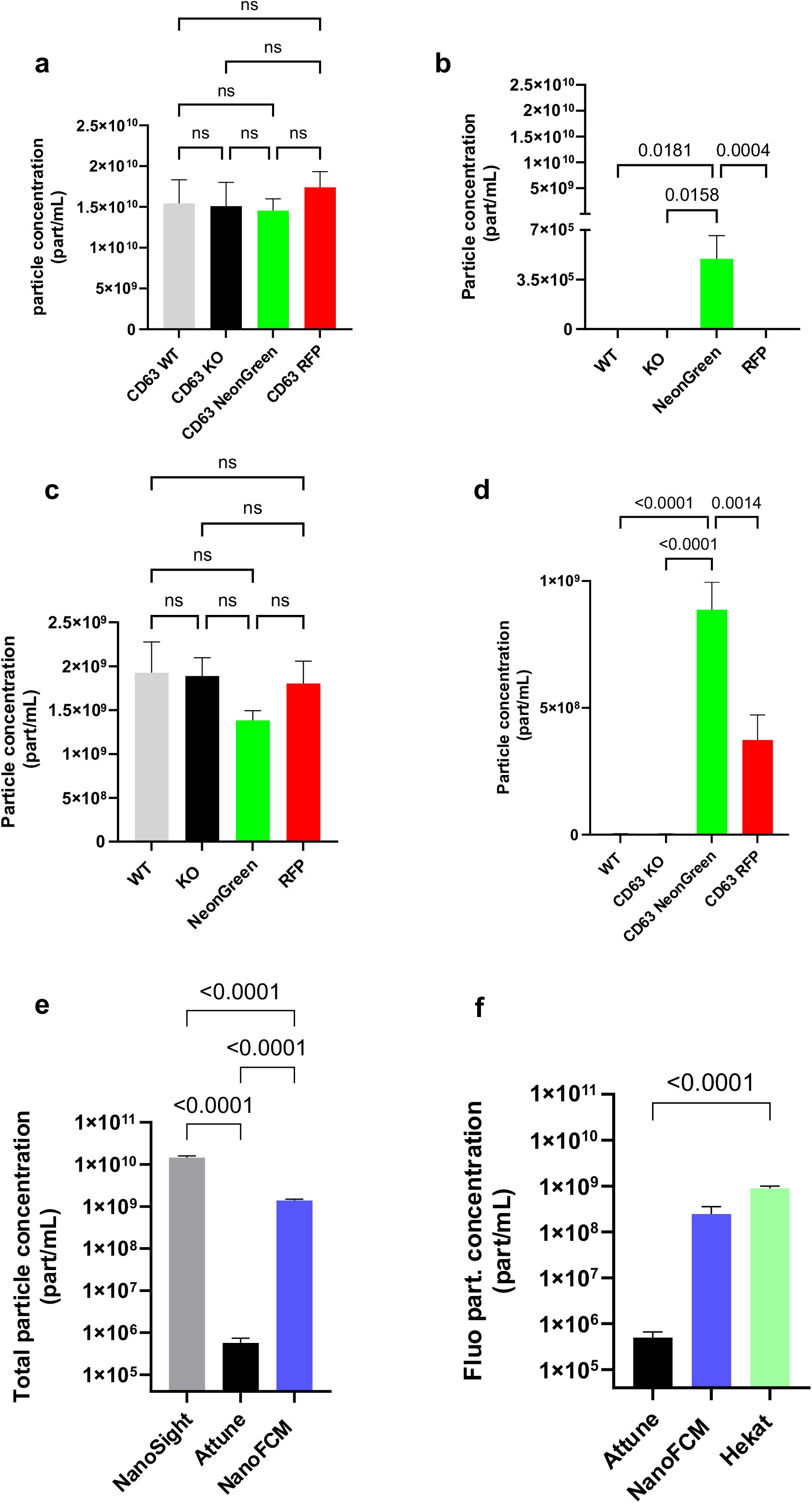
a. Particles concentration from HeLa WT, HeLa CD63 KO, HeLa mNeonGreen-CD63 and HeLa mRFP-CD63 derived sEV analyzed by NTA with NanoSight S300. HeLa WT and HeLa CD63 KO n=5, HeLa mNeonGreen-CD63 and HeLa mRFP-CD63 N=10. paired samples. Kruskal-Wallis test. **b.** Particles concentration from HeLa WT, HeLa CD63 KO, HeLa mNeonGreen-CD63 and HeLa mRFP-CD63 derived sEV analyzed with Attune NXT flow cytometer. Estimated with light scatter. HeLa WT and HeLa CD63 KO n=5, HeLa mNeonGreen-CD63 and HeLa mRFP-CD63 N=10. paired samples. Kruskal-Wallis test. **c.** Particles concentration from HeLa WT, HeLa CD63 KO, HeLa mNeonGreen-CD63 and HeLa mRFP-CD63 derived sEV analyzed with NanoFCM flow cytometer estimated with light scatter. HeLa WT and HeLa CD63 KO n=5, HeLa mNeonGreen-CD63 and HeLa mRFP-CD63 N=10. paired samples. Kruskal-Wallis test. **d.** Particles concentration from HeLa WT, HeLa CD63 KO, HeLa mNeonGreen-CD63 and HeLa mRFP-CD63 derived sEV analyzed with NanoSorter estimated by fluorescence detection. HeLa WT and HeLa CD63 KO n=5, HeLa mNeonGreen-CD63 and HeLa mRFP-CD63 N=10. paired samples. Kruskal-Wallis test. **e.** sEV concentration estimation from all devices by light scattering. Pooled measurements obtained during all acquisitions. NanoSight n=10, Attune NXT n=10, NanoFCM n=10. Paired samples. RM one-way Anova with Greisser-Greehouse correction test. **f.** sEV concentration estimation based on fluorescence detection with 488nm laser with three different devices. Attune NXT n=10, NanoFCM n=10, NanoSorter n=10. paired samples. Freidman test with Dunn’s multiple comparisons test.

We then proceeded to fluorescence-based detection to quantify fluorescently tagged particles, which is possible with flow cytometers and the NanoSorter. Using the Attune NXT flow cytometer, which is optimized for detecting EV subpopulations, we detected EV above background noise and identified a subset of mNeonGreen-positive particles (4.95 × 10⁵ ± 5.18 × 10⁵ particles/mL), confirmed to be associated with EV using Triton X-100 as a control, as recommended by the MIFlowCyt guidelines^25^ (Figure 2c and Supplementary Figure 2d). Although these results showed the presence of mNeonGreen-CD63 in EV, the estimated concentrations (1.7 × 10⁵ ± 3.7 × 10⁵ particles/mL) were lower than expected. nFCM, which is dedicated to nano-sized particles, was used to detect fluorescent particles. We observed a large population of mNeonGreen-positive EV (1.7 × 10⁹ ± 0.6 10^9^ particles/mL), which was confirmed by the absence of signal in the other conditions (Supplementary Figure 2e).

Unlike the previous instruments, the NanoSorter relies solely on fluorescence emission to detect particles. Thus, we observed large populations of mNeonGreen-CD63 (8.87 × 10⁸ ± 3.43 × 10⁸ particles/mL) and mRFP-CD63 (3.74 × 10⁸ ± 3.11 × 10⁸ particles/mL) particles, we were unable to detect non-fluorescent HeLa WT and HeLa CD63 KO EV. Notably, we detected a lower concentration of mRFP-CD63-positive EV compared to mNeonGreen-CD63 EV (Figure 2d). This difference may be attributed to variations in fluorescence intensity, stability, or expression levels per vesicle between the two fluorescent proteins.

Although each instrument relies on different technologies, they provided complementary insights into our EV batches, as we analyzed paired samples (Table 1). Total particle concentration based on light scattering revealed a much lower estimated concentration using standard flow cytometry compared to NTA and nFCM, even though these latter two also showed significant differences (Figure 2e). Fluorescence-based concentration estimates followed a similar trend, with standard flow cytometry detecting fewer fluorescent particles than nFCM and the NanoSorter. While standard flow cytometry provided a consistent ratio between scattered and fluorescent particles, its sensitivity may be insufficient to distinguish all EV from background noise. In contrast, nFCM and the NanoSorter, which are dedicated to nano-sized particles, detected high concentrations of fluorescent particles (Figure 2f).

**Table 1.**
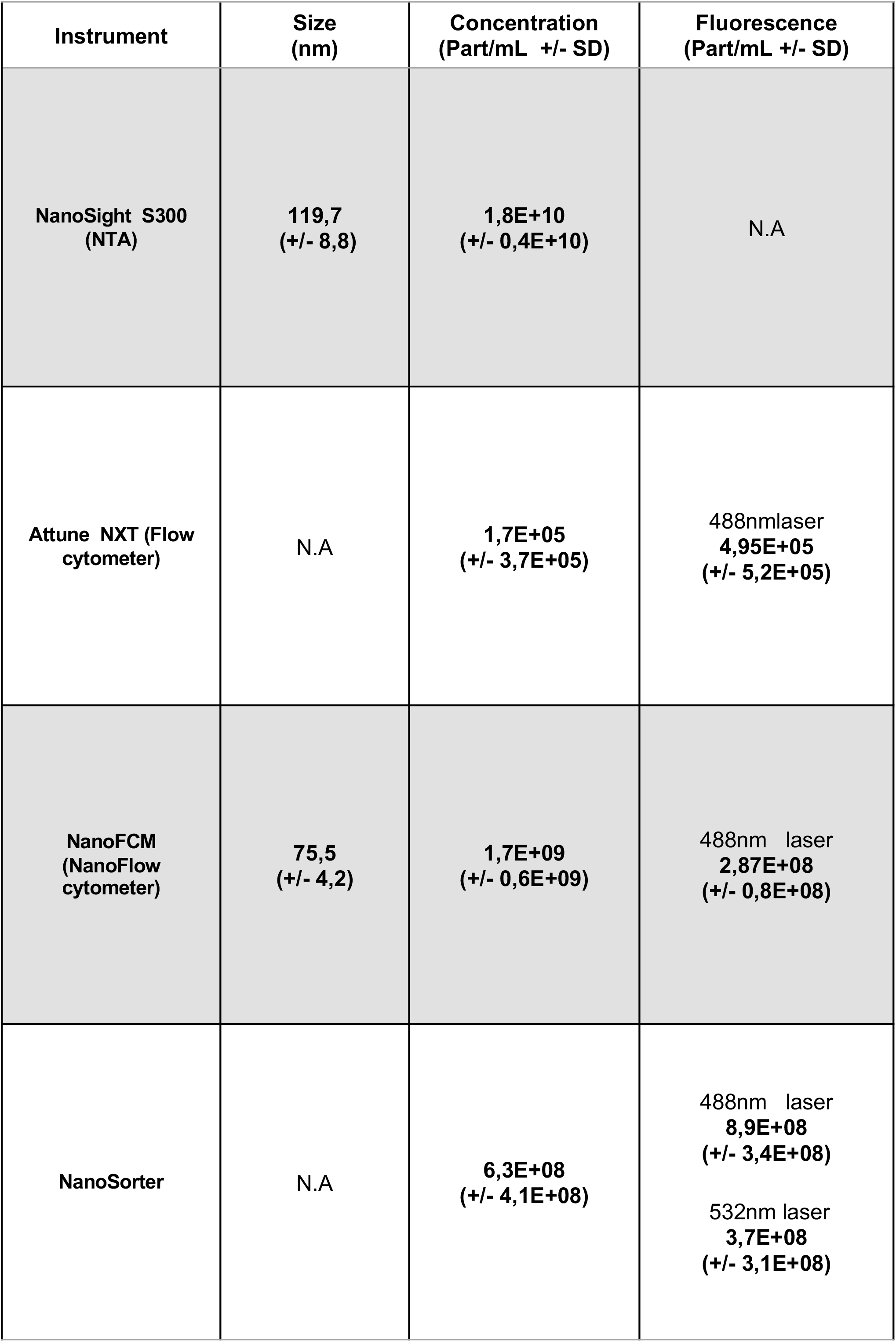

| Instrument | Size (nm) | Concentration (Part/mL +/- SD) | Fluorescence (Part/mL +/- SD) |
| --- | --- | --- | --- |
| NanoSight S300 (NTA) | 119,7 (+/- 8,8) | 1,8E+10 (+/- 0,4E+10) | N.A |
| Attune NXT (Flow cytometer) | N.A | 1,7E+05 (+/- 3,7E+05) | 488nmlaser<br>4,95E+05 (+/- 5,2E+05) |
| NanoFCM (NanoFlow cytometer) | 75,5 (+/- 4,2) | 1,7E+09 (+/- 0,6E+09) | 488nm laser<br>2,87E+08 (+/- 0,8E+08) |
| NanoSorter | N.A | 6,3E+08 (+/- 4,1E+08) | 488nm laser<br>8,9E+08 (+/- 3,4E+08)<br>532nm laser<br>3,7E+08 (+/- 3,1E+08) |

Thanks to these results, we confirmed the reproducibility of our EV production. The NanoSorter efficiently detected large amounts of fluorescent particles, comparable to instruments dedicated to nano-sized particle detection. However, the primary objective of the NanoSorter technology is to enable not only detection but also sorting and recovery of particles for potential downstream analysis.

### NanoSorter detects and sorts distinct particles with high specificity

The main feature of the instrument is its ability to encapsulate individual particles into droplets at high throughput and to selectively redirect these droplets after optical detection. This capability enables the recovery of sorted samples. Although similar functionality is available in conventional cell sorters, sorting extracellular vesicles (EV) remains challenging because of their small size, the limited detection sensitivity, and the large droplet volumes typically generated, making these systems complex to operate for EV applications. After sample loading, EV are encapsulated within aqueous droplets in the NanoSorter, using an oil-based sheath fluid. This process generates highly uniform droplets of approximately 40 fL, with diameters ranging from 4 to 5 µm (Figure 3a). In order to encapsulate single-EV into single droplets, we took advantage of the Poisson distribution^26^ to define the droplet size in order to obtain less than 3% of doublets at a theoretical concentration of 10^9^ particles/mL. mNeonGreen-CD63 EV and mRFP-CD63 EV isolated and normalized at the same concentration were mixed in two parallel experiments at a 70%/30% and 30%/70% ratio respectively. Detection by the NanoSorter of the mix before sorting revealed a proportion of the minor populations to be sorted between 26,67% for mRFP particles and 33,53% for mNeonGreen particles respectively. We then performed sorting of the minor populations that we analyzed first taking advantage of the microfluidic chips to keep the sorted particles within the droplets so we could observe them. Through fluorescent microscope, we could observe either mNeonGreen or mRFP fluorescence contained within droplets (Supplementary Figure 3a). In each case, whether we decided to sort mNeonGreen or mRFP particles, we observed by fluorescence microscopy high enrichment of the desired particles. These results suggested the ability of the NanoSorter to generate an enriched population of EV based on their fluorescence. To fully exploit the potential of single extracellular vesicle (EV) sorting for downstream analyses, we developed new strategies based on piezoelectric effect to merge droplets containing the EV of interest into a single larger droplet. We then recovered this 5fL droplet in 30μL of buffer to recover a maximum number of vesicles in a minimum volume (Supplementary Figure 3b). After 4 hours of sorting with an average sorting rate of 200 positive particles per second (200Hz), we could reach a population of approximately 4x 10^6^ EV finally recovered in a final volume of 30µL for each experiment. Downstream sorting of either mNeonGreen or mRFP CD63 EV minor population, detection measurements by the NanoSorter showed a clear enrichment of the minor EV population from 27% to 91% for mRFP-CD63 EV and from 34% to 89% for mNeonGreen-CD63 EV (Figure 3b).

**Figure 3.**
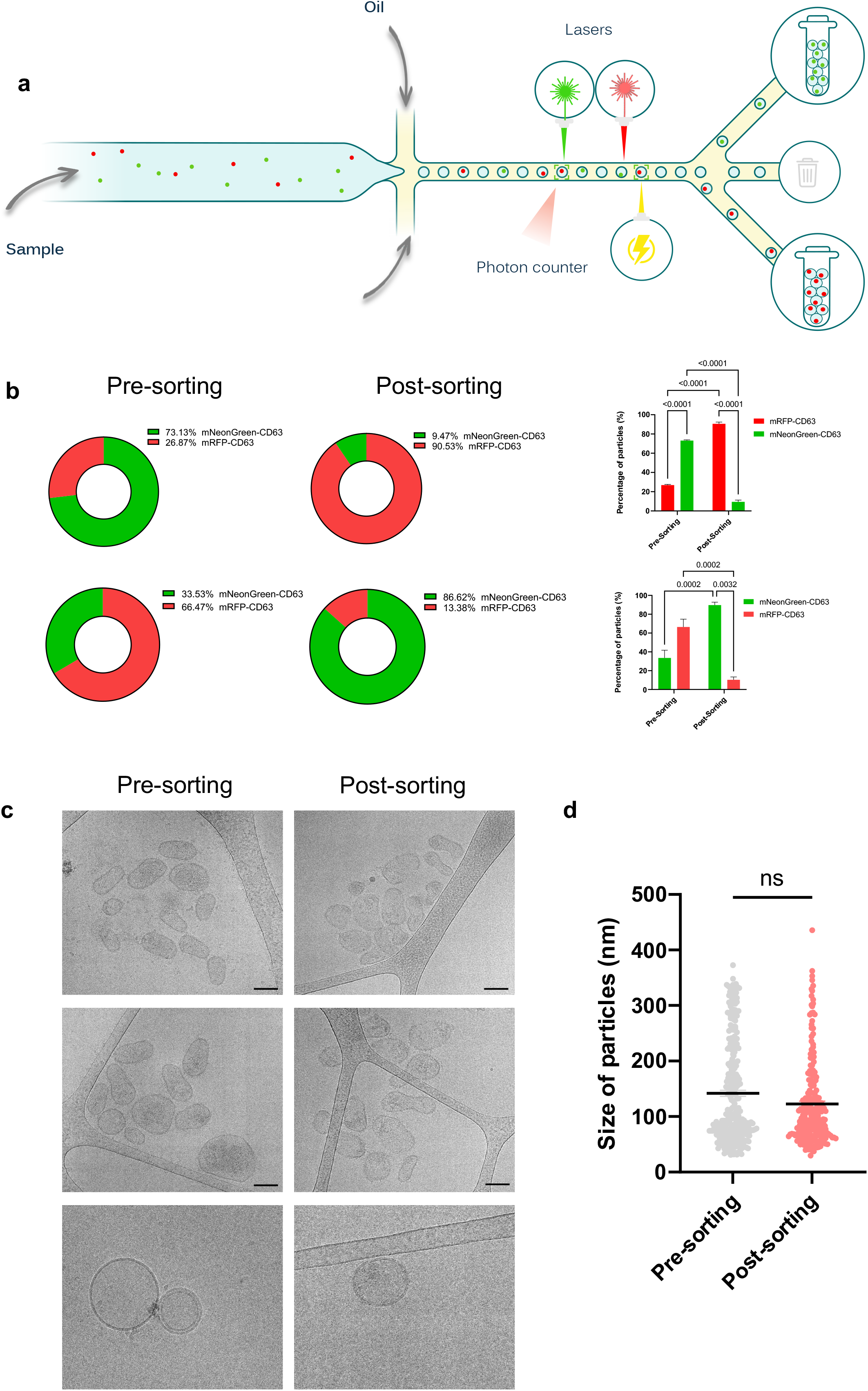
a. Scheme of EV processing through the NanoSorter for individual EV sorting. **b.** Measurement of mNeonGreen-CD63 or mRFP-CD63 positive EV population enrichment post sorting as part of whole and histograms before and post sorting. 3 independent experiments. 2-way Anova test. **c.** Tomograms of HeLa mNeonGreen-CD63 EV pre-sorting and post-sorting by cryo-EM. EV were processed through NanoSorter and recovered before being prepared for cryo-EM. Dark scale bar =100nm, white scale bar =50nm. 3 independent experiments. **d.** Size of particles measured by cryo-EM. Particles were measured in pre-sorting and post-sorting condition. Pre-sorting n= 292, post-sorting n= 228. Mann-Whitney test.

### NanoSorter sorting mode does not affect physical properties of sEV

The sorting workflow includes potential harming processes such as encapsulation of biological material, fluorescence detection through laser excitation of the fluorescent reporter, droplet deviation by dielectrophoresis and finally fusion of the sorted droplets. To ensure the integrity of sorted fraction and demonstrate the capacity of the device to sort a sufficient number of vesicles for downstream analysis, we investigated EV morphology by electron microscopy methods. Negative staining and Transmission Electron Microscopy (TEM) pictures of the original sample and the collected EV post-sorting revealed enough material to be analyzed. We did not observe any obvious change in morphology of the sorted HEK mNeonGreen-CD63 EV (Sup Figure 3c). Furthermore, the size of the sorted EV populations, measured on TEM pictures, did not change compared to the original sample (Sup Figure 3d). As EV processing for TEM includes chemical that affects their morphology^27^, we completed our observations by cryo-EM that preserves EV in a close-to-native state^28,29^. After 4h of sorting EV from HeLa cells stably expressing mNeonGreen-CD63, we collected 2.10^7^ particles of mNeonGreen-CD63 EV for each sample, that were prepared for cryo-EM imaging. We could observe that the integrity of the lipid bilayer was intact post-sorting and the morphology and the size of the EV were not altered (Figure 3d). Altogether, this showed that EV process by the NanoSorter did not alter their integrity and offered the possibility to harness EV post-sorting for downstream analysis.

### B lymphoma derived EV exogenous labelling allows sorting and enrichment of a specific EV subpopulation with a biomarker of interest

In order to extend the application of NanoSorter to the pathophysiological context and to explore its potential use in a clinical setting, we used antibody labeling approaches to detect potential biomarkers associated with EV. We targeted CD24, a highly glycosylated protein with a small protein core that is linked to the plasma membrane via a glycosyl-phosphatidylinositol anchor. CD24 is highly overexpressed in cancers^30,31^ and present on a subpopulation of EV^32^. We used two cell lines, an immortalized B cell line (BLCL) and the Namalwa cells, isolated from a patient diagnosed with Burkitt’s lymphoma, a particularly aggressive form of leukemia^33^. We determined by flow cytometry (Sup Figure 4a) and Western blot (Figure 4a) that CD24 was not detected in BLCL cells but was specifically expressed in Namalwa. This differential expression was even more striking on their respective EV (Figure 4a) providing a relevant model to test the NanoSorter. Analysis of EV by NTA revealed no changes in terms of particle concentration and size between the two cell lines (Figure 4b). To detect a potential enrichment of the CD24 signature to a sub-population of EV, we performed a first co-localization assay using anti-CD24 and tetraspanin mix (TMix) of anti-CD63, -CD9 and -CD81 antibodies on next generation NTA device possessing a fluorescent NTA mode (FI-NTA) and a colocalization system. We confirmed the association of CD24 with the Namalwa but not BLCL derived EV subpopulation (Figure 4c). Detection with nFCM revealed that CD24 was found in both CD9 or CD63 positive particles derived from B lymphoma cells (Sup Figure 4b). We confirmed the colocalization of CD24 with TMix in Namalwa but not BLCL derived EV with the NanoSorter, with a mean concentration estimated at 3,6.10^9 part/mL (Figure 4d) of double positive particles. Using NanoSorter’s technology, we sorted CD24/TMix double positive particles from a population of 2.10^9^ EV among which 6.10^8^ were fluorescent EV for 5hrs. By reinjecting the sorted samples containing from 3 to 5.10^6^ EV for detection, we observed a decrease of the Tmix single positive population by more than 60% while the double positive CD24+/Tmix+ population increased to 51,7% of the collected samples (Figure 4e). Even though we could observe the occurrence of unwanted events, a nearly 4-fold enrichment of double positive particles relative to the fluorescent EV populations was possible in our given conditions. Compared to the total EV populations, measured by NTA, the subpopulation of interest was enriched more than 8-fold in the post-sorting sample (Figure 4f). These data showed that the sorting capacity of the NanoSorter can be applied to endogenous EV after immunolabeling.

**Figure 4.**
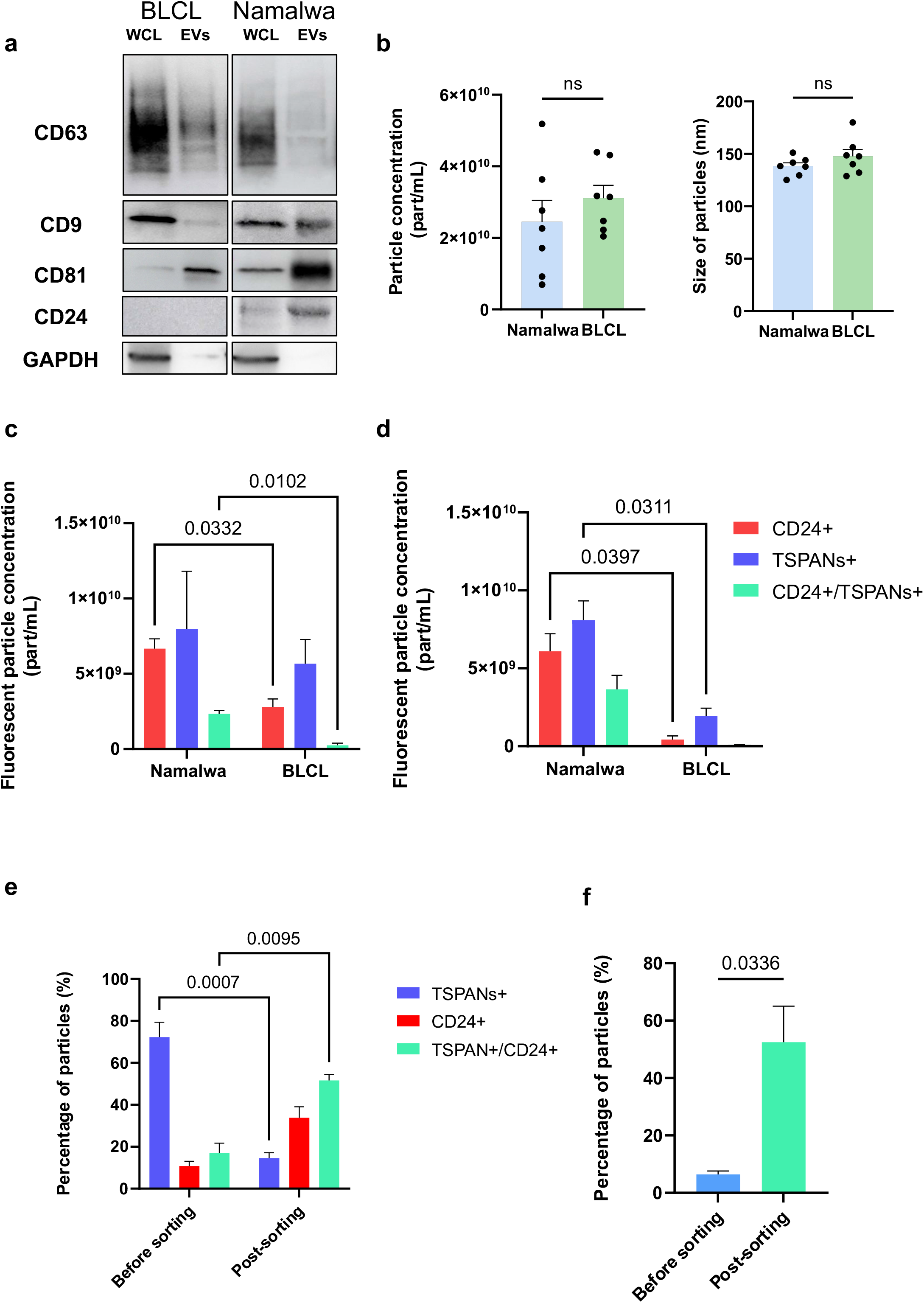
a. Western blot of Namalwa and BLCL whole cell lysates and derived EV isolated by SEC. 3 independent experiments. **b.** Size and concentration measurements of Namalwa derived EV and BLCL derived EV with Zetaview Evolution. Namalwa EV n=7, BLCL EV n=7. Student’s test with Welch’s correction. **c.** Concentration measurement of Namalwa derived EV and BLCL derived EV labelled with CD24 antibody and TSPANmix. Single positive population and double positive population were measured with Zetaview Evolution by FI-NTA and colocalization software. Namalwa EV n =3, BLCL EV n=3; 2-way Anova test. **d.** Concentration measurement of Namalwa derived EV and BLCL derived EV labelled with CD24 antibody and TSPANmix. Single positive population and double positive population were measured with NanoSorter. Namalwa EV n =4, BLCL EV n=5; 2-way Anova test. **e.** Measurement of fluorescent EV populations isolated from Namalwa derived EV labelled with CD24 antibody and TSPANmix before and after sorting with Zetaview. Single and double positive EV populations are shown as percentage of the total population measured. 4 independent experiments. 2-way Anova test. **f.** Measurement of fluorescent EV populations isolated from Namalwa derived EV labelled with CD24 antibody and TSPANmix before and after sorting among the total of particles measured by scattering with Zetaview. Double positive EV populations are shown as percentage of the total population measured. 4 independent experiments. Paired T-test.

## Discussion

Extracellular vesicles represent a potential subset of analytical biomarkers for research and clinical applications. Their composition, mirroring that of the producing cell contains valuable information, that help researchers to better understand the biology of the cell, its microenvironment and their contribution in cell-cell communication. Through isolation, detection and analysis, EV may become a major partner for disease research and diagnosis. Thanks to the next generation of isolation methods and analysis, many limits for EV research were pushed forward but remains associated with the investigation of bulk EVs. Here we assessed the ability of the NanoSorter to detect and most importantly to sort individual EV allowing the isolation of single EV sub-population for downstream analysis.

The NanoSorter detection and sorting capacities are based on EV associated fluorescence. We took advantage of the natural tetraspanins enrichment in EV to generate in vitro cell lines that produce fluorescent EV reporters on a CD63 CRISPR/Cas9 KO background. The localization of the rescued tetraspanins was comparable with endogenous WT proteins and was found in EVs without changing their main cargoes as described in the literature, suggesting that these reference EVs were relevant as biological samples. The stable overexpression allowed us to produce samples in a reproducible manner, allowing us to compare the results obtained with various detection devices. Although the expression of one fluorescent tetraspanin on EV does not reflect the diversity of vesicles present in biological fluids, it matched better the size, fluorescence and heterogeneity of biological samples than fluorescent silica or latex beads.

Thanks to our fluorescent reporters, we could assess this new technology which allows the detection of individual engineered fluorescent EV and to some extent, nano-biological objects, exogenously labeled, at a single particle level. The observed sensitivity of detection shows significant amounts of particles enabling the capacity to determine a concentration out of small samples thanks to the small flow of the device (10 µL/h) consuming a very small amount of sample. Compared to other devices able to quantify labeled particles, the NanoSorter analyzed a greater number of events acquired per analysis, providing greater statistical power, which enabled more robust and reliable data analysis. Importantly, the results are in good agreement with those obtained using commercially available instruments, demonstrating the validity of the approach.

Using the Hekat sorter, we could detect fewer mRFP-CD63 positive particles than mNeonGreen-CD63 EV. Since the detection sensitivity observed was comparable to NanoFCM with mNeonGreen-CD63 EV, the possibility that HeLa mRFP-CD63 cells either produce fewer fluorescent particles or sort less fluorescent protein per particle is not to be excluded and shall require further analysis. However, the detection and quantification of the particles are based on the fluorescence detection itself, so we overlook non-fluorescent particles or contaminants such as low density lipoparticles that are considered with other techniques based on indirect estimation like Brownian motion of particles. While the NanoSorter does give a precise estimation of a population of interest, this limitation may affect the adjustment of the parameters for efficient sorting. To prevent such underestimation and to adjust the droplet flow, a preliminary concentration estimation can be considered either with a complementary device or by using lipid dyes or a tetraspanin mix to cover a large spectrum of the EV subpopulation and limit non-specific sorting or doublets.

Importantly, the instrument can sort individual EV, resulting in the collection of a single EV population, out of the bulk sample without modifying the morphology of the particles as observed by cryo-EM. We showed here that the instrument allows the enrichment of specific fluorescent markers independently from the wavelength detected. Post-sorting, minor populations of fluorescent EV can be enriched more than 4 times and up to 8 times compared to the total of particles while other population of EV that may contaminate downstream analysis can be depleted by a factor of 7. The number of sorted EVs depends on various factors: the efficiency and relevance of the labeling, the relative abundance of the EV subpopulation within the biological samples and the limit of the NanoSorter. Yet, 4-5 hours of sorting can lead to the collection of 20 million EV, a quantity that would reach the requirements of downstream analysis as exemplified by cryo-EM morphological assessment. Importantly, observation of EV post sorting by cryo-EM did not reveal any change in their size, morphology or integrity. This provides an advantage for the NanoSorter compared to antibody-based isolation procedures of EV where the elution step can be detrimental to the vesicles and downstream analysis. However, we cannot conclude yet that the sorting process does not extract the protein corona that is loosely attached to the EV surface. Further studies will focus on complementary downstream analysis of the sorted samples, either by conducting refined characterization or functional assays in vitro or in vivo, to evaluate the functional integrity of EV post sorting.

Using CD24 as a theoretical EV-associated biomarker in the context of B lymphoma, we could first confirm its association with Namalwa-derived EVs. We could detect CD24 on CD9 positive EVs, in line with the major localization of both proteins at the plasma membrane of B cells. But a CD24 positive EV population, depleted for the three tetraspanins, the most widely used as EV reporters, was also detected. This highlights the heterogeneity of EV and the need to develop approaches to isolate EV subpopulations, which the NanoSorter is capable of. More importantly, we show that sorting of individual EV population based on exogenous labeling of two different markers. Although we used an artificial situation where CD24 is totally specific to cancer EVs, the Hekat sorter was able to enrich the subpopulation of EVs harboring this target. Considering the detection and sorting results with the NanoSorter, CD24 association with EVs may support its use as clinically informative material in this model of Burkitt’s lymphoma where it is overexpressed. As CD24 is also express by healthy cells during development or B cell maturation^34,35^, future studies will determine whether it may represent a biomarker of interest for Burkitt’s lymphoma. In any case, the selection of biomarkers associated with EVs in any pathology will require upstream optimization for optimal detection and sorting by the NanoSorter. Alternative therapeutic use of EV may also profit from the use of the NanoSorter. Detection of potential targets at the surface of EV may be beneficial for CAR-T cell therapy^36^ against this particularly aggressive form of leukemia that mostly affects children.

In conclusion, we describe here an instrument allowing sensitive detection of extracellular vesicles at a single EV level based on fluorescence. Associated with a cutting-edge microfluidic system, detection is followed by accurate sorting of single EV leading to the recovery of specific EV populations, based on at least one informative marker without altering their physical integrity. The NanoSorter is able to detect and sort EV originating from in vitro cell lines with transitory or stable expression of a fluorescent reporter. Moreover, the device is also prone to the detection of antibody labelled particles in purified samples. Finally, we show that the NanoSorter can detect specific biomarkers associated with a specific EV subset, which exhibits the potential interest of the instrument and EV in clinical situations for diagnostics or therapeutic follow-up.

## Methods

### Cell culture, transfection and drug treatments

HEK293 cells were obtained from ATCC. HeLa Kyoto cells were originally obtained from ATCC and given to us by B. Goud’s laboratory. HeLa CD63KO Cells were generated in the lab as previously described^21^. HEK293, HeLa and other cells were cultured in Dulbecco’s modified Eagle Medium (DMEM) Glutamax (Gibco) supplemented with 10% (vol/vol) fetal bovine serum (FBS) and 1% penicillin-streptomycin (Gibco). The Namalwa B lymphoma cell line isolated from a three-year-old patient with Burkitt’s lymphoma) and the immortalized B lymphocyte (BLCL K.EBV) from healthy donor were provided by Beatrice Clemenceau’s laboratory (INSERM U1307, CRCI²NA, Nantes, France). Namalwa and BLCL cell lines were cultured with synthetic media X-VIVO15 (Lonza) supplemented with 1% penicillin-streptomycin (Gibco). All cells were cultured at 37°C in a 5% CO2 incubator. For DNA transfection, cells were seeded at day 1 in T150 culture flasks and transfected with 5µg of plasmids using JetPrime Reagent at day 2 by following manufacturer’s instruction (Polyplus-Transfection).

### Reagent, Antibody, plasmids

#### Plasmids

pCMV-CD63-mRFPN1 (generated in the lab by Christing Bissig); pCMV-CD63-mNeonGreen (Kindly provided by Samir el Andaloussi’s lab); pLenti-CD63-mRFPN1, pLenti-CD63-mNeonGreen, psPAX2 (addgene 12260), pMD2.G (addgene 12259).

#### Primary antibodies

mouse Anti-CD63 (Clone H5C6, BD Bioscience 556019), mouse Anti-CD9 (Clone ML13, BD Biosciences 555370), mouse Anti-CD81 (Clone JS81, BD Bioscience 555675), rabbit Anti-RFP (Abcam ab62341) / rabbit Anti-mNeonGreen (Cell Signaling 41236S) / Anti-GAPDH (Invitrogen PA1-987) / rabbit Anti-Lamp1 ( Invitrogen, PA-1-654A) / Anti-Calnexin (Abcam ab10286) / Anti-Alix (Sigma SAB4200477) / Anti-Syntenin (Abcam ab19903) / Anti-CD24 (Clone ALB9, MA191384 Invitrogen) / mouse Anti-CD63-PE (BD Bioscience 557305) / Anti-CD63-AF488 ( Clone H5C6, Biolegend) / mouse Anti-CD81-PE (BD Bioscience 561957) / Anti-CD81-AF488 (Clone 5A6, Santa Cruz) / Anti-CD24-FITC (clone SN3 A5-2H10, Invitrogen) /Anti-CD24-PE (Clone SN3 A5-2H10, Invitrogen)/ Anti-CD24-APC (Clone SN3 A5-2H10, Invitrogen) / Anti-CD9-FITC (Clone ML13, Biolegend) / Anti-CD9-AF488 (Clone HI9A 10, BD) / Mouse IgG-PE ( MA1-10415, Invitrogen).

#### Secondary antibodies

anti-mouse HRP (Invitrogen 31430) / anti-rabbit HRP (Abcam ab6721) / anti-mouse AF488;647(Invitrogen A11029; A21236) / Anti-rabbit AF488;647 (Invitrogen A11008; A21245).

### Generation of stable cell lines

HeLa CD63 KO and CD9 KO cells were seeded in 24 well plates with complete media at 37°C prior to viral infection. Viral particles were produced by transfection of HEK 293 Lenti X cells with Lentiviral plasmid encoding protein of interests and plasmids encoding for envelop proteins plasmids for 24h. Conditioned media was harvested, filtered through a 0,45µm PES pore and stored at -80°C. HeLa CD63 KO cells were transduced with lenti-mRFP-CD63 or lenti-mNeonGreen-CD63 with a MOI of 10 for 24 hours at 37°C with Polybrene. 24h after infection the cells were selected with 1µg/mL puromycin for 48h.

### sEV production and isolation from conditioned culture media (CCM)

For isolation by Size Exclusion Chromatography (SEC), the conditioned media was centrifuged at 300g for 10mn at 4°C supernatant followed by 2,000g centrifugation for 25mn at 4°C. The supernatant was then concentrated by ultrafiltration (molecular weight cutoff (MWCO) of 10,000, Amicon 15 tubes, Merck-Millipore) to obtain concentrated conditioned medium. sEV were then isolated by SEC with qEV² Original 70nm columns on the Automatic Fraction Collector (IZON Science). Fractions 1 to 3 were collected and directly stored or pooled and concentrated again using a 10,000-MWCO filter (Amicon 4, Merck-Millipore). EV samples were used immediately after isolation or stored at −80 °C.

### Immunofluorescence microscopy

Cells were seeded on 1.5 glass coverslips in 24 well plates at 37°C, 5% CO2 with complete media. After any treatments or reaching 80% confluency, cells were fixed with 4% PFA in PBS diluted in cell culture media at 1:1 ratio (2% PFA final) for 15mn at RT. After 15mn Media containing PFA was removed and replaced by 2% PFA in PBS for 15mn at RT. PFA was then discarded and cells were washed and stored in PBS 1X. After Fixation steps, cells are quenched with 50mM Glycin in PBS for 20mn at RT followed by Permeabilization and blocking steps with 0.2% BSA, 0.2% Saponin in PBS for 30mn at RT. Cells were then incubated with primary antibodies diluted in 0.2% BSA, 0.2% Saponin in PBS for 30mn. After washing in BSA, Saponin PBS solution cells were incubated with secondary antibodies in blocking solution for 30mn at RT. Coverslips were finally mounted on superfrost slides with ImmuMount Mounting medium (Thermo).

Cells and EV samples were observed on a Nikon Ti-7 eclipse confocal microscopy with a 1.4 60X oil immersion objective and images were acquired using NIS-Element software. Post-acquisition analysis and quantifications were performed on FIJI software.

### Nano-flow cytometry analysis (NanoFCM)

NanoAnalyzer—NanoFCM, S/N FNAN30E22101948. Single-photon counting modules were used to simultaneously detect the side scatter (SSC) (using a bandpass filter: 488/10), green fluorescence (bandpass filter 525/40 The instrument was calibrated for particle concentration using 250 nm standard fluorescent silica beads (QC beads) and for particle sizing using a mixture of pre-mixed silica beads with diameters of 68, 91, 113 and 155 nm (S16M-Exo beads). Each EV sample was diluted with 1X PBS filtered through a 0.2 μm filter to reach a particle count within the optimal range of 2000–12,000 particles for 1 min at a pressure of 1 kPa. The recorded data was processed using the NanoFCM Profession 2.0 software to extract .fcs files

### Flow Cytometry Analysis

Cells were stained with anti-CD24-PE antibody (Clone SN3, Invitrogen) for 30mn at RT in PBS, and analyzed on a BD FACSymphony flow cytometer (BD Biosciences) using BD FACSDiva software. Unstained cells and cells stained with a matched isotype control antibody were used as negative controls to define gating boundaries and assess nonspecific binding. Data were analyzed with FlowJo version 10.10 software.

For EV, flow cytometry analysis was performed using Attune NXT flow cytometer with specific filter set. A threshold of 3000 ssc mvolt was applied to exclude background noise and aspecific events. SSC-H and FSC-H voltage were calibrated in order to distinguish buffer debris and sEV. Measurements were performed during 3 mn with at 12.5µL per minute debit. Acquisition of measurement and gating process including EV-Gate and mNeonGreen gate were performed using Attune NXT software. Post-acquisition Analysis was performed with FlowJow V10.10 software.

### Dynamic Light Scattering (DLS) Analysis

After isolation, sEV size and concentration were measured with Zetasizer Ultra ZSU5700 (Malvern Panalytical). Measurements of 50µL samples were performed with low-volume quartz batch ZEN 2112 cuvettes (Hellma Analytics, QS High Precision Cell, Art. No.105251006965-450, light path= 3×3 mm, Centre= 9.65) Cuvettes were cleaned with 70% EtOH, distilled water and dried prior to any measurements. Analysis of the samples was processed with ZS Xplorer v3.00 software (PSS0048-15) and considered as protein dispersed in water, at room temperature with a dispersant scattering value of 82.

Dynamic light scattering measurements of EV concentration were performed with a Zetasizer Ultra system (Malvern Panalytical) and ZS XPLORER (version 2.0.0.98) acquisition software. Samples were loaded into a low-volume quartz batch ZEN 2112 cuvette (Hellma Analytics, QS High Precision Cell, light path 3 × 3 mm) for analysis at room temperature with a dispersant scattering value of 82. The distributed particle concentration was measured.

### Nanoparticle Tracking Analysis (NTA)

EV samples size and concentration were measured on a NanoSight300 with a low volume flow cell gasket chip (NTA4169, Malvern Panalytical). After dilution, measurements were performed using standard reference EV protocol relying on 5 successive acquisitions of 1min before analysis and particle tracking. Mean size and concentration were obtained along with size distribution of population.

### Fluorescent Nanoparticle Tracking Analysis (FI-NTA)

Particle size and concentration were measured by NTA and fluorescence NTA (FI-NTA) using a ZetaView Evolution Quatt (Particle Metrix) equipped with. The instrument was calibrated before each session with 100 nm polystyrene beads for scatter mode, and with 488 nm and 640 nm monofluorescent beads for fluorescence mode. Measurements were performed using the “PMX PAN EV staining” predefined method, with gain 40, shutter 200, and frame rate 30 settings, at 25 °C and across 11 positions with 1 cycle per sample. Samples were diluted in PBS to reach 50-200 particles per frame. Total particle size and concentration were acquired in scatter mode, and the concentration of fluorescent particles was determined separately using the 488 nm and 640 nm lasers with the corresponding emission filters. Colocalization analysis was then performed to quantify double-positive particles (488⁺/640⁺) and to determine the fraction of particles carrying both fluorescent signals.

### Transmission Electron Microscopy (TEM)

For conventional EM, cells were seeded on coverslips and fixed with 2.5% glutaraldehyde in 0.1 M cacodylate buffer. The cells were processed for Epon (TAAB Laboratories Equipment) embedding as previously described^27^. Ultrathin sections were contrasted with uranyl acetate and lead citrate or Uranyless (Delta Microscopy).

For EV, samples were directly deposited on 200 mesh formvar/carbon-coated copper/palladium grids for 20 min at room temperature., fixed with PFA 2%/0.1 M phosphate buffer, and processed for immunogold labelling as for cells^27^.

### Cryo-Electron microscopy

Cryo-EM was performed on conventional. Briefly, 4,2 μL of sample was deposited on electron microscopy grids coated with a perforated carbon film. After draining the excess liquid with filter paper, grids were quickly plunged into liquid ethane and mounted onto a Gatan 626 cryoholder. Cryo-EM observation was performed with a TalosL120C (thermofischer) microscope operated at 120 kV, and images were recorded with a Ceta-M camera.

### Western blot

Cells were lysed on ice in a lysis buffer (20mM Tris, 150mM NaCl, 1% Triton X-100 pH 7.2) supplemented with a protease inhibitor cocktail (Roche). Cell lysates or EV were incubated with Laemmli sample buffer with or without 350mM 2-mercaptoethanol (Sigma), incubated at 95°C for 5 min, loaded on 4–20% Bis-Tris gels (Bio-Rad), and transferred on nitrocellulose membranes (Bio-Rad). The membranes were blocked in PBS/0.1% Tween (PBS/T) with 5% nonfat dried milk, incubated with primary and secondary antibodies diluted in PBS/T–milk. HRP signal was revealed using an ECL Clarity Max (Bio-Rad). Acquired bands intensities were quantified with ImageJ Fiji software (version 1.54f, Java 1.8.0_322 64 bit).

### Antibody Labelling of sEV

After isolation, EV samples were incubated with CD63-APC and CD24-FITC antibodies at 2.5µg/mL dilution for 30mn at RT in the dark. After incubation EV samples were washed with Exospin mini (Cell Guidance system) columns to remove unbound antibodies following manufacturer’s instructions. Eluate was then collected and processed for fluorescence detection and sorting with the NanoSorter.

### NanoSorter Analysis

Prior to analysis with the NanoSorter, EV samples could optionally be analyzed for total particle concentration. In this study, EV samples were analyzed using NTA prior to NanoSorter analysis to compare fluorescent particle counting with another instrument and to obtain an estimation of the total number of EVs.

EV samples were then loaded into counting chips for fluorescent detection at a precisely controlled flow rate of 5 or 10 µL/h. Post analysis, when the sample concentration was too high to ensure the detection of individual particles within the detection zone, the sample was diluted to below 5×10^9^ particles per ml

### NanoSorter Sorting experiments

Fluorescent EV samples with a concentration below 5 × 10⁹ particles/mL could be injected into the microfluidic sorting chip, either for single-color or two-color sorting. Sorting was performed at frequencies ranging from 3.5 to 5 x10^4^ droplet per second.

For widefield microscopy observations, sorted droplets containing fluorescent EVs were injected into a counting chip and placed under microscope equipped with 488nm or 520nm lasers with a beam expander. Images were captured with the Kinetix camera C-mounted on the microscope.

For downstream applications, the droplets containing objects of interest were collected in a sheath fluid trap directly connected to the collection channel of the sorting chip. Then, the trap was then secured, and the droplets were fused by applying piezoelectric impulses. All droplets containing objects of interest were then fused into a single droplet. This droplet was injected into a microtube for analysis or store, with an additional volume of medium added when necessary to facilitate liquid handling.

### Statistical analysis

Statistical tests were performed using GraphPadPrism 10.6.1 (892) version.

### Image analysis

Image J Fiji (version 1.54f, Java 1.8.0_322 64bit) was used for microscopy and western blot analysis. FlowJow V10.10 was used for Flow Cytometry analysis.

### Statistics and reproducibility

All graphs with statistical tests represent the quantification of at least 3 independent experiments. Significant p-value are shown as absolute value. Graphs represent the means of quantification for each experimental condition with SEM.

## Acknowledgement

The authors acknowledge the Cytocell - Flow Cytometry and FACS core facilty (SFR Bonamy, BioCore, Inserm UMS 016, CNRS UAR 3556, Nantes, France) for its technical expertise and help, member of the Scientific Interest Group (GIS) Biogenouest and the Labex IGO program supported by the French National Research Agency (n°ANR-11-LABX-0016-01). We acknowledge the MicroPICell core facility (SFR Bonamy, BioCore, Inserm UMS 016, CNRS UAR 3556, Nantes, France), member of the Scientific Interest Group (GIS) Biogenouest, IBISA, and the national infrastructure France-Bioimaging supported by the French national research agency (ANR-24-INBS-0005 FBI BIOGEN). We acknowledge the CBMN and Sisareuth TAN and Dr. Olivier LAMBERT for their technical expertise and help for cryo-TEM observations. The authors are Thankful to Alan Adamczyk for Flow cytometry help and advice.

## Author Contribution

MC wrote the manuscript, conceptualize the study and performed most experiments (lead); EL performed experiments (minor); LBC & JCD performed analysis (minor); KS & CR performed NanoSorter development and analysis; PG & SB NanoSorter development and funding (lead); BC provides essential reagents and feedback; MG Nanosorter development and analysis, supervision of the detection and sorting with the NanoSorter, review manuscript (lead); GVN supervision and conceptualization of the study, review manuscript, secured funding experiments (lead).

## Ethical

Generation of GMO was conducted under approval from French Governmental Research and Innovation General Direction trough the approval n°11374.

## Supplementary Data

**Supp Figure 1.**
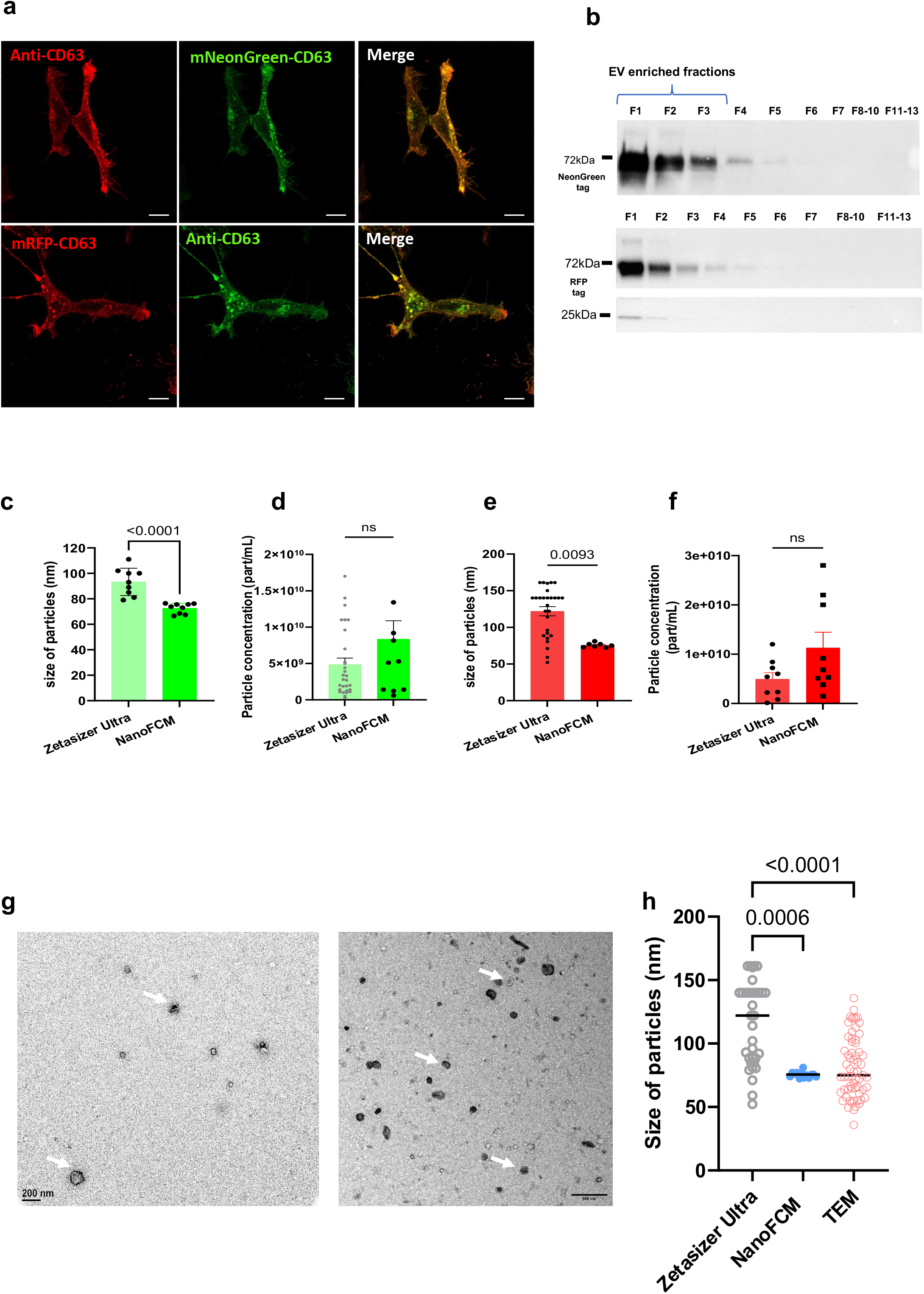
**a.** localization of fluorescent reporter in HEK WT with mNeonGreen-CD63 transitory expression and HEK WT with mRFP-CD63 transitory expression. Cells were labelled for Lamp1. Endogenous fluorescence for both fusion proteins were observed. Scale bar 10um. 3 independent experiments. **b.** Western blot of size exclusion chromatography (SEC) fractions from HEK WT with mNeonGreen-CD63 transitory expression and HEK WT with mRFP-CD63 transitory expression. Fractions 8 to 10 and 11 to 13 were pooled before loading into SDS-PAGE gel wells. Samples were immunoblotted for their respective fluorescent tag along with CD63 and CD9. 3 independent experiments. **c.** Size of HEK WT with mNeonGreen-CD63 transitory expression derived sEV isolated by SEC. Particle size was measured either by DLS with Zetasizer Ultra or with NanoFCM. 3 independent experiments. n= 9, Mann-Whitney test. **d.** Particle concentration of HEK WT with mNeonGreen-CD63 transitory expression derived sEV isolated by SEC. Estimated concentration was measured either by DLS with Zetasizer Ultra or with NanoFCM. 3 independent experiments. n= 9, Unpaired t-test. **e.** Size of HEK WT with mRFP-CD63 transitory expression derived sEV isolated by SEC. Particle size was measured either by DLS with Zetasizer Ultra or with NanoFCM. 3 independent experiments. Zetasizer n= 18, NanoFCM n = 9, Kruskal-Wallis test. **f.** Particle concentration of HEK WT with mRFP-CD63 transitory expression derived sEV isolated by SEC. Estimated concentration was measured either by DLS with Zetasizer Ultra or with NanoFCM. 3 independent experiments. Zetasizer Ultra n=18, NanoFCM n= 11, Kruskal-Wallis test. **g.** Tomograms of HEK mNeonGreen-CD63 and mRFP-CD63 transient expression derived EV isolated by SEC. Isolated EV were processed for transmission electron microscopy. Scale bar 100nm. 3 independent experiments. **h.** Size of isolated EV from HEK with both transient expressions. Diameter of particle was measured from TEM observations and compared to size estimation with Zetasizer Ultra and NanoFCM. Zetasizer Ultra n=34, NanoFCM n=13, TEM n=64. Kruskal-Wallis test.

**Supp Figure 2.**
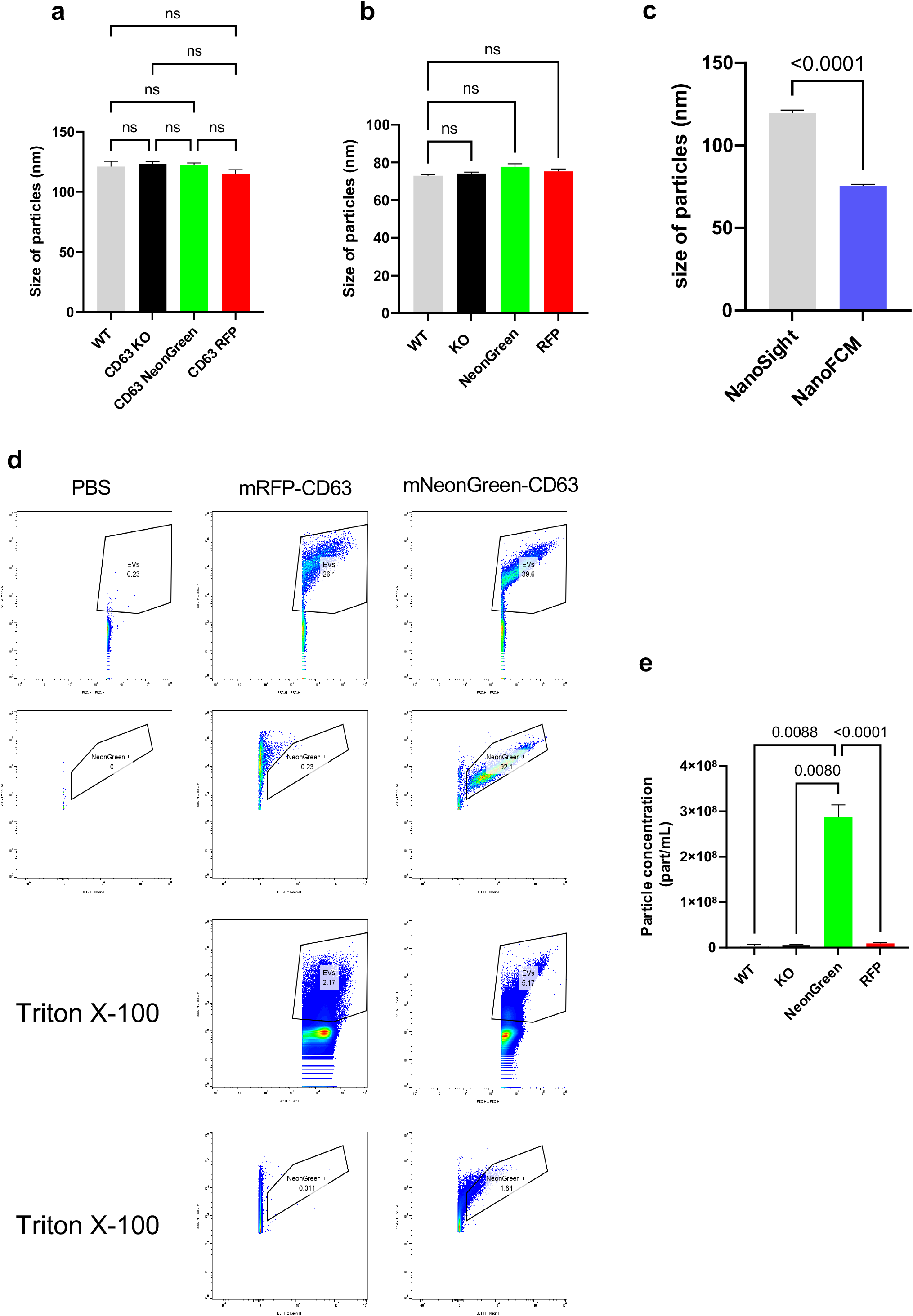
**a.** Size distribution of EV derived from HeLa WT, HeLa CD63 KO, HeLa mNeonGreen-CD63 and HeLa mRFP-CD63 stable cell lines. Measurements were performed by NTA with NanoSight S300. Histogram of quantification of particle size for each condition. HeLa WT n=5, HeLa CD63 KO n=5, HeLa mNeonGreeN-CD63 n=10, HeLa mRFP-CD63 n=10. Ordinary One-way Anova test. **b.** Particle size of EV derived from HeLa WT, HeLa CD63 KO, HeLa mNeonGreen-CD63 and HeLa mRFP-CD63 stable cell lines. Measurement were performed with NanoFCM flow cytometer. Histogram of quantification of particle size for each condition. HeLa WT n=5, HeLa CD63 KO n=5, HeLa mNeonGreeN-CD63 n=10, HeLa mRFP-CD63 n=10. Kruskal-Wallis test. **c.** Particle size of EV derived from HeLa WT, HeLa CD63 KO, HeLa mNeonGreen-CD63 and HeLa mRFP-CD63 stable cell lines pooled. Measurements were performed by flow cytometry with NanoFlow Analyzer and by NTA with NanoSight S300. NanoSight n=30, NanoFCM n=30. Paired samples, Wilcoxon test. **d.** Representative dot plots of of EV derived from HeLa WT, HeLa CD63 KO, HeLa mNeonGreen-CD63 and HeLa mRFP-CD63 stable cell lines. Measurements were performed by flow cytometry with ATTUNE NXT. EV gating was performed with FSC and SSC followed by mNeonGreen detection with 488nm laser to generate NeonGreen+ gate. Dot plots of representatives gatings along with PBS as and mNeonGreen-CD63 EV treated with Triton X-100 as controls. HeLa WT n=5, HeLa CD63 KO n=5, HeLa mNeonGreeN-CD63 n=10, HeLa mRFP-CD63 n=10. **e.** Particle concentration of mNeonGreen EV derived from HeLa WT, HeLa CD63 KO, HeLa mNeonGreen-CD63 and HeLa mRFP-CD63 stable cell lines. Measurements were performed by flow cytometry with NanoFlow Analyzer and 488nm laser. HeLa WT n=5, HeLa CD63 KO n=5, HeLa mNeonGreeN-CD63 n=10, HeLa mRFP-CD63 n=10). One-way Anova test.

**Supp Figure 3.**
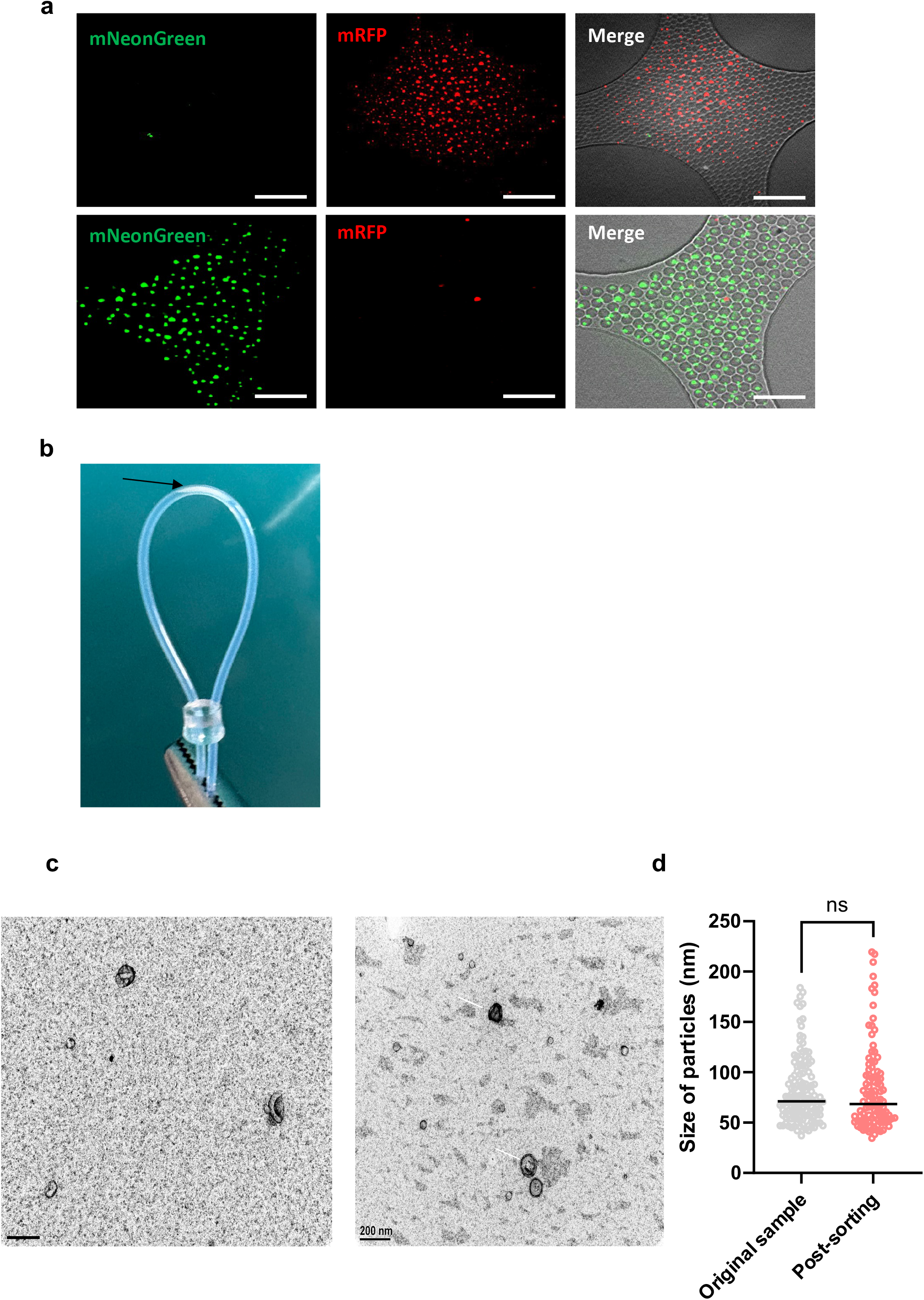
**a.** Observation of HEK mNeonGreen-CD63 or HEK mRFP-CD63 derived EV post sorting within droplets through fluorescent microscope. Histogram of positive mNeonGreen or mRFP particles after sorting either mNeonGreen or mRFP particles. **b.** Illustrations of droplets collection post sorting with NanoSorter. Illustrate droplets gathering (dark arrow) through microfluidic loop. **c.** Tomograms of HEK WT with mNeonGreen-CD63 transitory expression before and after sorting with the NanoSorter by Transmission Electron Microscopy. Scale bar 200nm. **d.** Quantification of particle size by TEM observations between original and post sorting samples (Original sample n=137, Post-sorting sample n=126). Mann-Whitney test.

**Supp Figure 4.**
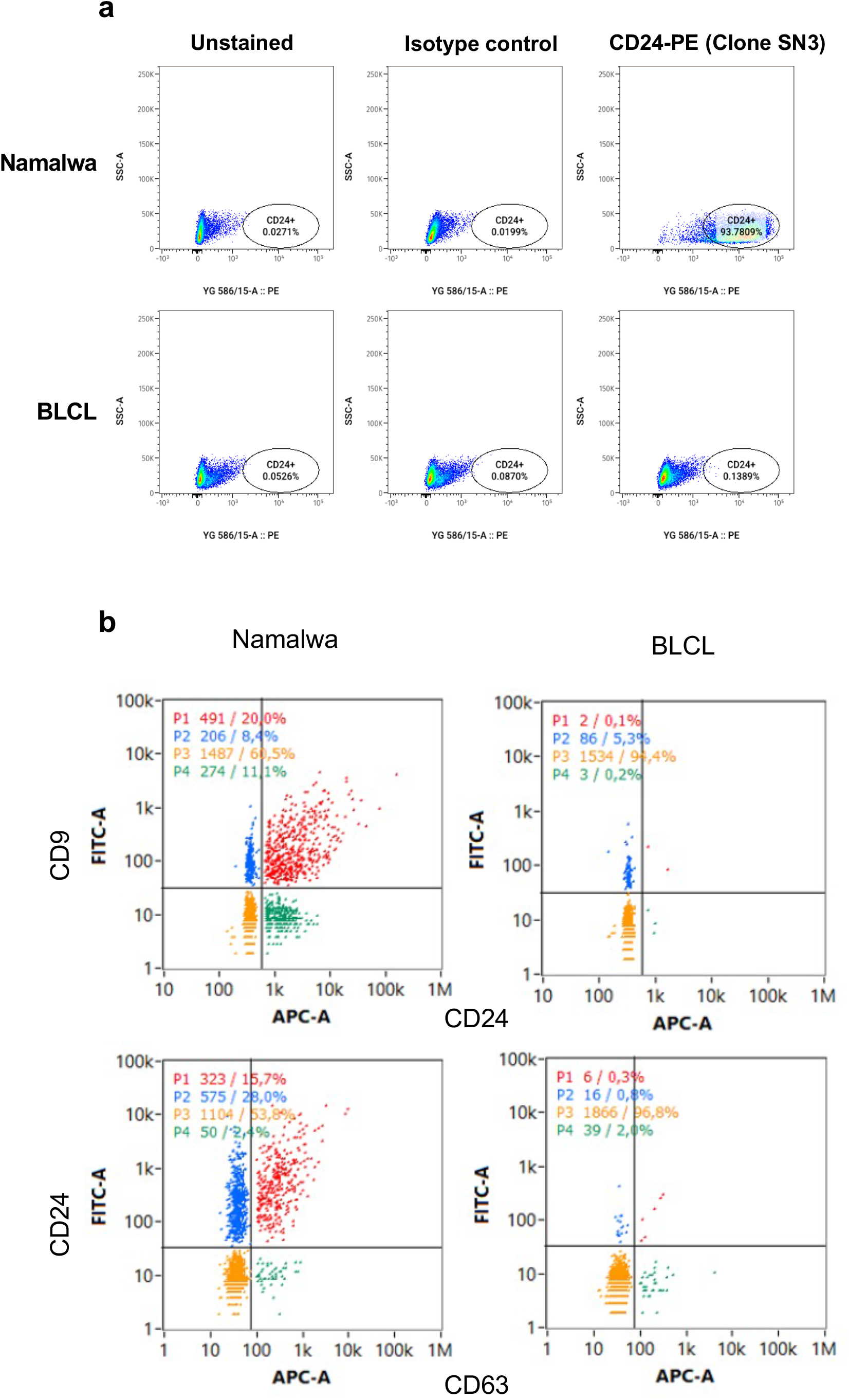
**a.** Labelling and detection of CD24 in Namalwa or BLCL cell lines by flow Cytometry with BD Symphony flow cytometer. Cells were gated with FSC and SSC to exclude debris and doublets. CD24-PE gate was then created to select CD24 positive cells. Mouse IgG-PE was used as isotype control. As shown representative dot plots of CD24-PE in unlabeled, isotype and CD24 labelled conditions. 3 independents experiments. **b.** Labelling of Namalwa and BLCL derived EV. EV were incubated with CD24-PE or FITC and Either CD63 or CD9 antibodies. Detection was performed with NanoFlow Analyzer. Dot plots as representative of each labelled population for Namalwa and BLCL derived EV. for each double positive population. 3 independent experiments.

